# The evolution of Hutchinsonian climatic niche hypervolumes in gymnosperms

**DOI:** 10.1101/2025.01.26.634928

**Authors:** Fernanda S. Caron, David F. R. P. Burslem, Juliano Morimoto

**Affiliations:** Programa de Pós-graduação em Ecologia e Conservação, Universidade Federal do Paraná, Curitiba, 82590-300, Brazil; School of Biological Sciences, University of Aberdeen, Cruickshank Building, Aberdeen AB24 3UU, UK; Institute of Mathematics, University of Aberdeen, King’s College, Aberdeen AB24 3FX, UK

**Author notes:** **Corresponding author:** Juliano Morimoto, Institute of Mathematics, University of Aberdeen, AB24 3UE, Aberdeen. **Data availability statement** Raw data is available through DOI links in supplementary material as the files were too large to attach with the submission, while code is available in supplementary material [during the review process to safeguard anonymity] and the Github repository: xxxxxx.

**Keywords:** climatic niche, persistence homology, phylogenetic comparative data, phylogenetic signal

## Abstract

**Aim:** The niche is a fundamental concept in theoretical and experimental ecology and is used to describe a wide range of ecological processes from species interaction with the environment to community assemblies. A common way to represent the niche is through a multidimensional geometry known as the Hutchinsonian niche hypervolume. Ecological theory predicts that niche hypervolumes have properties such as holes with broader eco-evolutionary significance, but we lack a comprehensive empirical study of niche hypervolume properties and their evolutionary significance.

**Location:** Global

**Time period:** Holocene

**Major taxa studied:** Gymnosperms

**Methods:** We conducted for the first time a systematic and comprehensive test of the evolution of Hutchinsonian niche hypervolume properties (volume and holes) across 65 genera and 12 families of gymnosperms, which includes many species that are endangered or threatened. Using cutting-edge computational algorithms, we measured the evolution of geometric (i.e. volume) and topological (i.e. holes) properties of gymnosperm hypervolumes across a comprehensive calibrated phylogeny.

**Results:** Our comparative analysis revealed weak evidence of the non-independent evolution of niche hypervolume volume and no evidence of the non-independent evolution of hypervolume holes. We also found that genera and families with low hypervolume volume such as monotypic groups like *Gingko*, likely experienced shifts in hypervolume evolutionary rates.

**Main conclusions:** Our results show that geometric and topological properties of gymnosperm climatic niche hypervolumes evolve independently. This agrees with competitive exclusion hypothesis in ecological theory where extant groups are likely to be the ones which minimise niche overlap and competition.

## INTRODUCTION

The ‘niche’ is arguably one of the most important concepts in ecology (Chase, 2011; McCann & Gellner, 2020). Although definitions are debated (Hutchinson, 1957; Soberón, 2007; Wennekes et al., 2012), the concept of the niche has enabled unprecedented theoretical and empirical advances in our understanding of how species interact with their environment (e.g., Blonder, 2016; Soberón & Nakamura, 2009; Soberón, 2010; Winemiller et al., 2015; and references therein). In particular, the definition of niche formulated by Hutchinson – the Hutchinsonian niche hypervolume – is widespread in the literature and is simple to apply to the large quantities of ecological data that are increasingly available. In this context, the niche hypervolume can be defined as “an abstract mapping of population dynamics onto an environmental space, the axes of which are abiotic and biotic factors that influence birth and death rates” (verbatim from Holt, 2009; see also Hutchinson, 1957). Niche hypervolumes have had a major influence on the advancement of ecological theory (Soberón & Arroyo-Peña, 2017; Blonder, 2018; Jiménez et al., 2019; Mammola & Cardoso, 2020; Soberón & Peterson, 2020; Carrasco et al., 2022; Conceição & Morimoto, 2022), as well as to uncover insights into niche evolution (Blonder, 2018; Pili et al., 2020; Bates & Bertelsmeier, 2021), niche differentiation (Carvalho & Cardoso, 2020; Huang et al., 2024), biological invasion (Tingley et al., 2014; Helsen et al., 2020; Zhang et al., 2020), community diversity (Chrisholm & Pacala, 2010; Loke & Chrisholm, 2023), ecological specialisation (Bebber & Chaloner, 2022), and global biodiversity patterns (Beaugrand et al, 2020; but see Justus, 2019).

The niche hypervolume concept has evolved with advances in theoretical and empirical ecology (Chase & Leibhold, 2003; Holt, 2009; Wiens, 2011; Kearney et al., 2017; Letten et al., 2017). One aspect of the hypervolume concept that has recently emerged, but remains untested, is the idea that hypervolumes can have holes (Blonder, 2016). Specifically, Blonder (2016) proposed the existence of holes in niche hypervolumes and conjectured that these holes could indicate environmental conditions in which species are vulnerable to be outcompeted in interspecific competition or unaccounted eco-evolutionary processes (e.g. range shifts) that lead to vacant niches. One cannot rule out the possibility of missing data generating artefacts interpreted as holes, but it is plausible – although largely untested in large scales – that holes have ecological meaning (Blonder, 2016; Conceição and Morimoto, 2022). Note that, in this conjecture, holes are representative of a fundamental ecological feature that could evolve if, for example, sister lineages share the physiological or morphological vulnerabilities that facilitate invasion portions of environmental space by other lineages. Blonder (2016) tested the holey niche concept in the morphological niche hypervolume of a community of co-existing finches in the Galapagos and concluded that “well-known datasets may be described in terms of hypervolumes with holey geometries that are potentially consistent with unexplored mechanisms” (Blonder, 2016). Thus, Blonder’s conjecture could link niche hypervolume properties that exist in high-dimensional space to real-world ecological processes, making the study of hypervolume properties a paramount surrogate and key target of investigation to understanding species’ ecology. However, to date, we still lack large scale tests to directly ascertain the evolution of Hutchinsonian climatic niche hypervolume geometric and topological properties. Yet, the concept of climatic niche hypervolumes was originally created to represent species’ climatic occupancy and it what hypervolumes are widely used in many applications today (Blonder, 2018; Vilas et al., 2022).

Studying niche hypervolume properties is not trivial. The incomplete information available for defining the dimensions of the hypervolume topology poses a challenge. Patchy and heterogeneous observations prevent reconstruction hypervolume topology from bioclimatic variables with accuracy (e.g., Peterson et al., 2018; Jiménez et al., 2019), and statistical models to “fill” hypervolume topologies are criticized (e.g., Qiao et al., 2017; Mammola, 2019; Guillerme et al., 2020). These limitations can be particularly insidious when estimating hypervolume holes as high dimensionality can pose a challenge (Blonder, 2016; Conceição and Morimoto, 2022). We recently proposed the use of algebraic topology, specifically the concept of persistence homology, to identify holes in hypervolumes and overcome the dimensionality challenges (Conceição & Morimoto, 2022). However, neither we nor others have yet used approaches to identify and derive biological meaning from holes and other properties from ecological niche hypervolumes across species within an evolutionary framework. Such comparative studies are needed to better understand if and how ecological niche hypervolume properties, including holes, evolved.

In this paper, we addressed an important knowledge gap about niche hypervolumes, namely, if niche hypervolume properties evolve non-independently in related species. To achieve this, we present the first comparative study of the evolution of niche hypervolume properties (volume and holes). Understanding evolutionary patterns of niche hypervolume properties can be a way to uncover the eco-evolutionary significance of its properties such as holes, which have been hypothesised to hold meaningful ecological information but still lack large-scale studies to test this. Plants are excellent models for studying hypervolume properties because they minimise potential confounding effects of range shifts and migration observed in animals (e.g., Valladares et al., 2014; Menchetti et al., 2019; Häfker et al., 2022; and references therein). Many plants are threatened by climate change and understanding the evolution of niche hypervolumes may uncover new insights into how to protect them (Antão et al., 2020; Singh et al., 2023). Moreover, climate niche evolution in plants is representative of similar processes in animals (Liu et al., 2020).

Here, we focused our attention on studying ecological niche hypervolume properties of gymnosperms. Gymnosperms non-flowering woody plants with major importance for many ecosystems worldwide. Gymnosperm trees are abundant in many subtropical, temperate, and boreal forests that collectively represent over one-third of the forested regions on the planet (Lesiv et al., 2022). Moreover, gymnosperms form the group of plants has assessed as having high extinction risks (∼40% of its species classified as CR, EN, VU) despite their importance for cultural and provisioning ecosystem services (Forest et al., 2018; Tagliari et al., 2023), whereas other gymnosperms are a primary source of timber (e.g., *Pinus sp*). Gymnosperms are also relatively less diverse than flowering plants, likely due to direct competition (Bond, 1989; Coiffard et al., 2012; De Boer et al., 2012), which makes them a tractable but representative group for investigating the evolution of ecological niche hypervolume properties. Given gymnosperms’ broader evolutionary and socio-ecological significance, we analysed the environmental niches of 65 gymnosperm genera distributed across 12 families (Figure 1B). We used genera and families, rather than species, as our taxonomic levels of comparison because the immense computational power to estimate hypervolume geometric and topological properties prevents a species-level analysis at this stage, even for high-performance computers. For example, calculating hypervolume holes in a single hypervolume with more than a few thousand of data points in dimensions >2 is virtually impossible with current algorithms, and we still lack a properly validated approach to rarefy the point cloud (i.e. reduce number of points) without losing topological information. We constructed niche hypervolumes with 31 environmental variables and applied a recently developed approach of topological data analysis to estimate hypervolume holes (Conceição & Morimoto, 2022). We hypothesised that if niche hypervolume properties captured conserved ecological processes, then hypervolume properties would display strong phylogenetic signals. For example, if closely related genera and families are more likely to have more similar climatic niches (e.g. over- or under-dispersed point clouds in niche space), then this would be captured by our comparative analysis of climatic niche properties such as volume. Our findings advance our conceptual and biological understanding of the multidimensional niche and the evolution of its properties.

**Figure 1.**
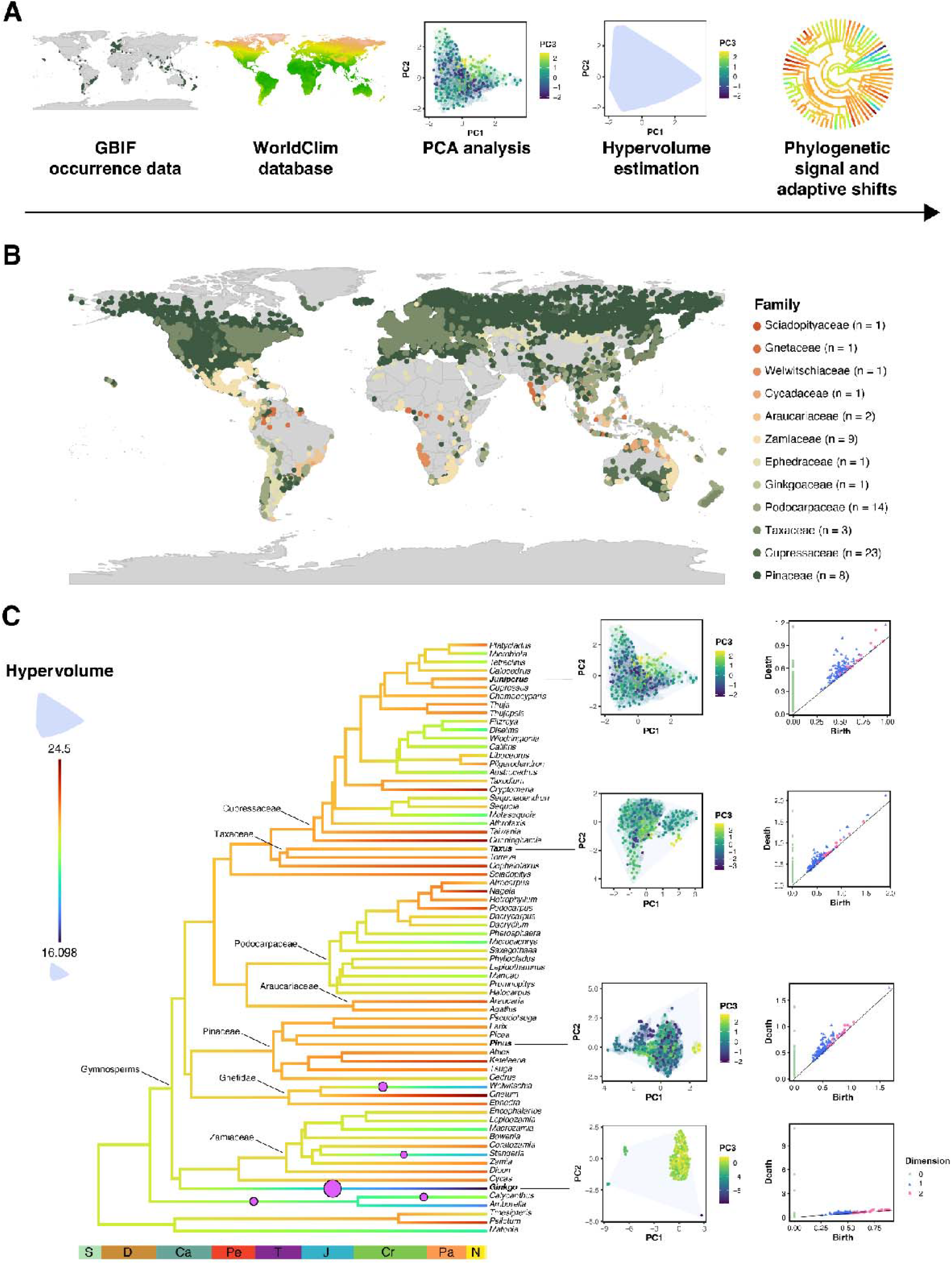
Overview of the analyses performed in the present study. **A**. Steps to analyse the data: collection of the occurrence data in GBIF database; extraction of the bioclimatic variables for each occurrence; PCA analysis with the bioclimatic data; estimation of the climatic niche hypervolume volume and properties; calculation of the phylogenetic signal of the hypervolume properties. **B**. Occurrence data for each family studied. Legend indicates the number of genera in each family. **C**. Hypervolume properties, phylogenetic signal, and adaptive shifts estimated. Phylogeny is mapped with hypervolume volume, in which blue corresponds to small hypervolume volume and red to large volumes. Time scale is represented at the base of the phylogeny. Families represented by more than one genus are highlighted in the internal branches. Circles represent the adaptive shifts with a posterior probability higher than 0.3. Graphs at the right represent examples of hypervolumes estimated and the persistence homology diagrams. Green: dimension 0, Blue = dimension 1, and Pink: dimension 2 homology.

## MATERIALS AND METHODS

### Data collection

We obtained occurrence data from the GBIF database (https://www.gbif.org/) for all available gymnosperms. The search was conducted for Cycadopsida (40,554 entries), Ginkgoopsida (57,488 entries), Pinopsida (5,723,563 entries), and Gnetopsida (52,648 entries) (Table S1). We processed the GBIF data to remove duplicated points, points with locations in impossible places (e.g., the sea), and points with large uncertainty. We assumed that observations outside native ranges were valid in the construction of the realised niche hypervolume and were therefore maintained. In this respect, we interpreted the realised climatic hypervolume niche as the multidimensional representation of all measured climatic conditions where the genus or families have survived (either within its native range or naturalised in an alien area). Recent studies have used hypervolumes to test the differences in climatic niche in invasive vs native ranges (e.g. Tingley et al. 2014; Zhang et al., 2020) but this is not the scope of this paper. We only kept points derived from human observation, as opposed to fossil records or preserved specimens. The final dataset contained 2,483,316 entries, corresponding to 764 species. The taxonomic classification for the gymnosperms was followed according to the World of Flora Plant List (http://www.worldfloraonline.org/, last accessed Oct 13, 2023).

Apart from the gymnosperm data, we also gathered occurrence data for four outgroup families (see Data analyses section). These families were Psilotaceae (23,865 entries), Calycanthaceae (10,013 entries), Amborellaceae (327 entries), and Matoniaceae (836 entries) (Table S1). For these families, we had a total of 16,361 entries after cleaning and processing the data, corresponding to 23 species. Climatic data were obtained from the WorldClim database using the getData function in raster v3.6-26 (Hijmans et al., 2023). This dataset was downloaded at a resolution of 2.5 minutes of a degree, using the argument “var” as “bio” and “tmin”. Phylogenetic relationships for the genera and families for which we had ecological data were retrieved from Stull et al. (2021). In this paper we only considered the climatic component of the niche, as climatic variables represent the primary determinant of variation in plant distributions globally (e.g. Walter, 1979). We acknowledge that numerous additional abiotic and biotic factors contribute to niche diversification in plants, including soil properties, elevation and topography, but incorporating these niche dimensions is beyond the scope of the current paper.

### Data analyses

We performed analyses of the evolution of the Hutchinsonian niche across gymnosperms at the level of genera and families. We discuss the limitations of this approach in the discussion section. We used 31 environmental variables to construct the climatic niche of each taxon. First, we extracted the climatic variables related to temperature and precipitation from the WORLDCLIM database (https://www.worldclim.org/) with resolution of 5 minutes for the occurrence points of each taxon using the extract function in *raster* v3.6-26 (Hijmans et al., 2023). Then, we conducted a principal component analysis (PCA) with all climatic variables to reduce the dimensionality of these environmental variables to three orthogonal axes (i.e., PC1-3). A PCA was conducted for each gymnosperm genus and family separately using the function prcomp() in R (R Core Team, 2024). For each taxon, we also calculated their centroid in the first three PC axes. For this, we calculated the mean of each PC axis for the taxa individually and for the gymnosperms. Then, we estimated the Euclidean distance between each taxon centroid to the gymnosperm centroid. While the hypervolume volume is based on the extreme values of the PCA, the distance between centroids can be regarded as a measure of the uniqueness of the taxon’s climatic niche. Hypervolumes were estimated using the hypervolume_gaussian function in *hypervolume* v.3.1-4 (Blonder, 2024) with default parameters. This algorithm provides random points that “fill” the hypervolume space (see Blonder et al., 2014) and from which properties of the hypervolumes can be estimated. As the hypervolume analyses are computationally intensive, we had to reduce the dataset of the genera and families that had more than 50,000 locality entries. The genera that had their occurrences reduced were *Abies, Juniperus, Larix, Picea, Pinus*, and *Taxus*, and the families were Cupressaceae and Pinaceae. Inspection of the occurrence points derived from the entire and reduced dataset showed that this reduction in the locality entries did not affect the coverage of the taxon occurrence (Figure S1). Finally, we estimated the phylogenetic signal and adaptive shifts for the hypervolume properties and the distance of centroids. The phylogenetic signal was calculated using the phylosig function in *phytools* v2.3-0 (Revell, 2024). The statistics used were Pagel’s λ (Pagel, 1999), with 1,000 iterations, and Blomberg’s *K* (Blomberg et al., 2003), with 1,000 simulations. We also mapped the hypervolume properties in the phylogeny using the contMap function in *phytools* v2.3-0 (Revell, 2024). As a final analysis, we estimated adaptive shifts in the evolution of the hypervolume volume using the bayou package v2.3-0 (Uyeda & Harmon, 2014). This package fits Bayesian reversible-jump multi-optima Ornstein–Uhlenbeck (OU) models to phylogenetic comparative data, identifying the location and magnitude of adaptive shifts (Uyeda & Harmon, 2014). We ran the analyses at the genus and family levels using the default prior configurations. We ran the models three times at each level for 10 million generations, considering 30% as burn-in. All variables were log-transformed prior to the calculations. The phylogeny used was manipulated to be used at the genus and family level, keeping either one random representative per genus or family for this. In addition, we also estimated the dimension 2 holes in the 3D ecological hypervolumes using a state-of-the-art topological data analysis approach (Conceição & Morimoto, 2022) and the *TDAstats* package (Wadwa et al., 2018). We focused on dimension 2 holes because they represent the boundaries of 3D sub-volumes within the 3D hypervolume. We estimated maximum niche hypervolume hole size (i.e. largest hole size) as the maximum “survival” of dimension 2 holes upon Vietoris-Rips filtration and the average and standard error as the mean and standard error of the survival of all dimension 2 holes identified for a given hypervolume (Conceição & Morimoto, 2022). Our motivation for studying these holes was that, if holes exist in ecological niche hypervolumes and evolve non-independently, we could infer their biological significance from the shared ecological properties among species. We did not focus on dimension 0 (holes which emerge based on distance among data points) or dimension 1 (holes in the faces of the hypervolumes) because they are dependent on the distribution of datapoints which defines the hypervolume rather than holes in the structure of our 3D hypervolumes (see Conceição & Morimoto, 2022 and references therein). It is important to note that we cannot rule out that hypervolumes constructed using principal components could distort the holes that exist thereby adding noise to our phylogenetic analysis (false negative). We currently lack a systematic study to ascertain whether our approach, which is common practice in ecology, conserves the topological structures of the niche hypervolume. Such knowledge will have far-reaching implications for ecological studies as many studies rely on the ability of computational algorithms to estimate the topology of niche hypervolumes accurately. All analyses were performed in R v4.4-0 (R Core Team, 2024).

## RESULTS

We first tested whether ecological niche hypervolume volume evolved non-independently across genera of gymnosperms. We found weak evidence to support the non-independent evolution of ecological niche hypervolume volume as only Blomberg’s *K* at the genus level was statistically significant (Table 1). Estimates of Pagel’s λ at the genus and family levels and Blomberg’s *K* at the family level were not statistically significant (Table 1 and Table S2).

**Table 1.** Phylogenetic signal using Blomberg’s K (Blomberg et al., 2003) and Pagel’s □ (Pagel, 1999) of hypervolume volume and holes at the genus level. LogL (logL0) refers to the log-likelihood estimates of the (null) models.

| Trait | $\lambda$ | logL | logL0 | p | K | p |
| --- | --- | --- | --- | --- | --- | --- |
| Hypervolume | 0.817 | -128.686 | -129.517 | 0.197 | 0.606 | **0.009** |
| Mean distance | 1.000 | 5.322 | 5.322 | 1.000 | 0.425 | 0.358 |
| Maximum distance | 0.000 | -24.736 | -24.736 | 1.000 | 0.397 | 0.503 |
| SD distance | 0.000 | -15.040 | -15.039 | 1.000 | 0.378 | 0.653 |

Next, to gain further insight into the weak support for the non-independent evolution of hypervolume volume at the genus level, we measured adaptive shifts in evolutionary rates for the hypervolume volume. Comparing the value of the adaptive regime (θ) in the root of the phylogeny with the adaptive regimes leading to other lineages, it is possible to assess if these new adaptive regimes suffer an increase in value or decrease. Thus, this can indicate if a smaller or larger hypervolume volume has been favoured in these lineages regarding their ancestor. In gymnosperms, the root value of the adaptive regime (θ) of the hypervolume volume was estimated as 21.548. Our model showed five major shifts in the adaptive regimes at the genus level leading to the evolution of smaller hypervolume volumes (Figure 1C). The clades which went through these decreasing shifts encompassed (1) *Welwitschia* (θ = 18.614, PP [posterior probability] = 0.438), (2) *Stangeria* (θ = 18.752, PP = 0.355), (3) *Gingko* (θ = 17.273, PP = 0.839), and the outgroup (4) *Calycanthus* (θ = 18.282, PP = 0.310), and (5) *Calycanthus* and *Amborella* (θ = 19.169, PP = 0.361). These results were virtually identical across all three runs and thus, are robust.

We also analysed the evolution of hypervolume holes (Blonder, 2016). However, none of our analyses of the average size of dimension 2 holes (Figures S2A, S3B), standard error of estimates of dimension 2 holes (Figures S2B, S3C), and maximum size of dimension 2 holes (Figures S2C, S3D) showed evidence of non-independent evolution of hypervolume holes. And while the overall shape and associated persistence diagrams differed amongst themselves (Figure 1C), pairwise distance matrices of the dimension 2 holes did not reveal any clusters of pairwise distances which appeared similar (Figures S2E, S3F). In fact, there was no difference between the observed and randomised pairwise distance matrices (Figure S4).

Lastly, we tested if the relative distinctiveness of the niche of each genus and family of gymnosperms evolved non-independently, but the results were not statistically significant (Text S1). These results suggest that ecological niche hypervolume holes, like hypervolume volume, likely capture more dynamic ecological processes that do not necessarily evolve non-independently among lineages.

## DISCUSSION

All species have a niche, and their niche are often represented as niche hypervolumes. Thus, the niche hypervolume remains one of the most significant concepts in ecology today (Chase, 2011; McCann & Gellner, 2020). Despite its significance to theoretical and empirical ecology, we lacked a comprehensive understanding of hypervolume properties and whether they evolve non-independently in related species. Here, we answer this question by studying the evolution of the Hutchinsonian climatic niche hypervolume properties in gymnosperms. We mapped how climatic niche hypervolume geometrical and topological properties, such as volume and holes, evolved across the gymnosperms. Our data supported the idea that climatic niche hypervolume properties evolved independently across lineages. There was no phylogenetic signal on hypervolume volume except for a statistically significant estimate of Blomberg’s *K* at the genus but not family levels. Blomberg’s *K* captures the degree of variation within and among lineages and a statistically significant value of *K* suggests that within lineage variation supersedes variation among lineages (Meireles et al., 2020). The statistically significant results at the genus but not family level were driven by the relatively homogenous hypervolume volumes of genera within the Pinaceae, Araucariaceae, and Podocarpaceae families (Figure 1B). These are genera and families with widespread climatic tolerances which are represented by hypervolumes with comparatively higher volumes in our analysis. Contrastingly, Pagel’s λ is a scalar that correlates a trait matrix with a phylogenetic matrix and was not statistically significant in any of our estimates. This is not surprising given the variability of hypervolume volume across gymnosperms found in our data both at the genus and family levels (Meireles et al., 2020).

We found a nearly 2-fold difference between the smallest and largest hypervolume by volume in our dataset. *Gingko, Stangeria*, and *Welwitschia* genera had the lowest hypervolume volume estimates (Figure 1C). *Ginkgo* is a living fossil that has been deemed one of the most critically endangered gymnosperms in the world and, therefore, has a relatively narrow ecological distribution reflected in its hypervolume volume (Forest et al., 2018). Although not classified as (critically) endangered by the IUCN Red List, *Welwitschia* has a unique ecology and occupies arid and semi-arid environments in sub-Saharan Africa with relatively narrow ecological distribution, also being considered a living fossil. Likewise, *Stangeria* is not classified as (critically) endangered by the Red List but has a unique ecology and is endemic to southern Africa (Schönland, 1918). These findings confirm that our hypervolume approach captures ecological characteristics of the genera and families and, thus, that the lack of evidence for the non-independent evolution of ecological niche hypervolume properties is meaningful. The causes of such shifts are unknown and multifactorial, but the direct competition and diversification of angiosperms likely contributed, although cannot fully explain this pattern. Adaptive shift results at the family level are available in Text S1.

Multidimensional hypervolumes are complex high-dimensional geometries which poses challenges for visualisation and computations of its properties. To study climatic niche hypervolume properties across gymnosperms, we needed to make simplifying assumptions to guarantee the feasibility of the analysis. One such assumption was that PCA transformation and dimensionality reduction of our point cloud preserved ecological information. This assumption is shared with most studies using the concept of niche hypervolumes (e.g., Pianka et al., 2017; Zhang et al. 2020; Ellis, 2022) but it is worth highlighting that no systematic study has yet tested the implications of PCA on point cloud structure and hypervolume properties (but see also Mahoney et al., 2017; Lu et al. 2021). PCA acts as a projection of the point cloud into lower dimensions and can introduce biases such as for example “filling in” holes which are present in higher dimensions. Similar biases can occur from our decision to measure hypervolume properties at the genus and family levels, since holes which are present in the climatic niche hypervolume of individual species might be filled in when analysing the climatic niche hypervolume of the genus or family. However, it is important to highlight that the opposite is not true: we cannot have artificial holes or topological structure emerging after PCA or at the genus or family levels if these structures are absent in the raw, species-level data. Thus, our study provides a conservative estimate of the evolution of climatic niche hypervolume properties. We are currently working to develop algorithms which directly test the influence of PCA on climatic niche hypervolume properties, and which would allow the computation of hypervolume properties (particularly holes) more efficiently. This will be the scope of future work.

## Conclusion

The niche is a cornerstone concept in ecology which has stimulated unprecedented advances in theoretical and empirical studies. Hutchinsonian climatic niche hypervolume is widespread in the literature and likely to become increasingly common for modelling ecological niches from open data. We studied the properties of the climatic niche hypervolumes of gymnosperms and found evidence that the climatic niche hypervolume properties evolve independently in gymnosperms. These are the first results to systematically probe how Hutchinsonian hypervolumes properties evolved, and to show that ecological niche hypervolume properties likely capture dynamic ecological characteristics of the niche. This probably limits the range of evolutionary inferences which can be made from Hutchinsonian climatic niche hypervolumes. Future work in other taxa for which phylogenetic relationships have been well established (e.g., vertebrates) will provide further insights into the evolutionary insights that can be gained from the concept of the niche hypervolume.

## REFERENCES

Antão, L.H., Weigel, B., Strona, G., Hällfors, M., Kaarlejärvi, E., Dallas, T., Opedal, Ø.H., Heliölä, J., Henttonen, H., Huitu, O. and Korpimäki, E., (2022). Climate change reshuffles northern species within their niches. Nature Climate Change, 12(6), pp.587–592.

Bates, O. K., & Bertelsmeier, C. (2021). Climatic niche shifts in introduced species. Current Biology, 31(19), R1252–R1266.

Beaugrand, G., Kirby, R. & Goberville, E. (2020). The mathematical influence on global pat-terns of biodiversity. Ecology and Evolution, 10, 6494–6511.

Bebber, D. P., & Chaloner, T. M. (2022). Specialists, generalists and the shape of the ecological niche in fungi. The New Phytologist, 234(2), 345.

Blomberg, S. P., Garland Jr, T., & Ives, A. R. (2003). Testing for phylogenetic signal in comparative data: behavioral traits are more labile. Evolution, 57(4), 717–745.

Blonder, B. (2016). Do Hypervolumes Have Holes? The American Naturalist 187. PMID: 27028084, E93–E105.

Blonder, B. (2018). Hypervolume concepts in niche-and trait-based ecology. Ecography, 41, 1441–1455.

Blonder, B. (2024). hypervolume: High Dimensional Geometry, Set Operations, Projection, and Inference Using Kernel Density Estimation, Support Vector Machines, and Convex Hulls. R Package Version 3. 1–4. https://CRAN.R-project.org/package=hypervolume

Blonder, B., Lamanna, C., Violle, C. & Enquist, B. J. (2014). The n-dimensional hypervolume. Global Ecology and Biogeography, 23, 595–609.

Bond, W. J. (1989). The tortoise and the hare: ecology of angiosperm dominance and gymnosperm persistence. Biological Journal of the Linnean Society, 36(3), 227–249.

Carrasco, J., Lisón, F., Jiménez, L., & Weintraub, A. (2022). A new method to estimate the ecological niche through n-dimensional hypervolumes that combines convex hulls and elliptical envelopes. bioRxiv, 2022-03.

Carvalho, J. C. & Cardoso, P. (2020). Decomposing the causes for niche differentiation between species using hypervolumes. Frontiers in Ecology and Evolution, 8, 243.

Chase, J. M. (2011). Ecological niche theory. The theory of ecology, 93–107.

Chase, J. M., & Leibold, M. A. (2003). Ecological niches: linking classical and contemporary approaches. University of Chicago Press.

Chisholm, R. A., & Pacala, S. W. (2010). Niche and neutral models predict asymptotically equivalent species abundance distributions in high-diversity ecological communities. Proceedings of the National Academy of Sciences, 107(36), 15821–15825.

Coiffard, C., Gomez, B., Daviero-Gomez, V. & Dilcher, D. L. (2012). Rise to dominance of angiosperm pioneers in European Cretaceous environments. Proceedings of the National Academy of Sciences of the United States of America 109, 20955–20959.

Conceição, P. & Morimoto, J. (2022). ‘Holey’niche! finding holes in niche hypervolumes using persistence homology. Journal of Mathematical Biology, 84, 58.

De Boer, H. J., Eppinga, M. B., Wassen, M. J., & Dekker, S. C. (2012). A critical transition in leaf evolution facilitated the Cretaceous angiosperm revolution. Nature communications, 3(1), 1221.

Ellis, C. J. (2022). A hypervolume approach to niche specialism, tested for the old-growth indicator status of calicioids. The Lichenologist, 54(6), 379–387.

Forest, F., Moat, J., Baloch, E., Brummitt, N.A., Bachman, S.P., Ickert-Bond, S., Hollingsworth, P.M., Liston, A., Little, D.P., Mathews, S. and Rai, H., (2018). Gymnosperms on the EDGE. Scientific reports, 8(1), p.6053.

Guillerme, T., Puttick, M. N., Marcy, A. E., & Weisbecker, V. (2020). Shifting spaces: Which disparity or dissimilarity measurement best summarize occupancy in multidimensional spaces?. Ecology and evolution, 10(14), 7261–7275.

Häfker, N. S. et al. (2022). Animal behavior is central in shaping the realized diel light niche. Communications Biology 5, 562.

Helsen, K. et al. (2020). Inter-and intraspecific trait variation shape multidimensional trait overlap between two plant invaders and the invaded communities. Oikos, 129, 677–688.

Hijmans, R. J., van Etten, J., Sumner, M., Cheng, J. Baston, D., Bevan, A., … & Wueest, R. (2023). raster: Geographic Data Analysis and Modeling. R Package Version 3. 6–26. https://CRAN.R-project.org/package=raster

Holt, R. D. (2009). Bringing the Hutchinsonian niche into the 21st century: ecological and evolutionary perspectives. Proceedings of the National Academy of Sciences 106, 19659–19665.

Huang, J., Yu, R., Ding, Y., Xu, Y., Yao, J., & Zang, R. (2024). The Relationship between Trait-Based Functional Niche Hypervolume and Community Phylogenetic Structures of Typical Forests across Different Climatic Zones in China. Forests, 15(6), 954.

Hutchinson, G. E. (1957). Population studies-animal ecology and demography-concluding re-marks in Cold Spring Harbor symposia on quantitative biology, 22, 415–427.

Jiménez, L., Soberón, J., Christen, J. A., & Soto, D. (2019). On the problem of modeling a fundamental niche from occurrence data. Ecological Modelling, 397, 74–83.

Justus, J. (2019). Ecological theory and the superfluous niche. philosophical topics, 47, 105–124.

Kearney, M., Simpson, S. J., Raubenheimer, D., & Helmuth, B. (2010). Modelling the ecological niche from functional traits. Philosophical Transactions of the Royal Society B: Biological Sciences, 365(1557), 3469–3483.

Lesiv, M., Schepaschenko, D., Buchhorn, M., See, L., Dürauer, M., Georgieva, I., … & Fritz, S. (2022). Global forest management data for 2015 at a 100 m resolution. Scientific Data, 9(1), 199.

Letten, A. D., Ke, P. J., & Fukami, T. (2017). Linking modern coexistence theory and contemporary niche theory. Ecological Monographs, 87(2), 161–177.

Liu, H., Ye, Q., & Wiens, J. J. (2020). Climatic-niche evolution follows similar rules in plants and animals. Nature Ecology & Evolution, 4(5), 753–763.

Lu, M., Winner, K., & Jetz, W. (2021). A unifying framework for quantifying and comparing nDdimensional hypervolumes. Methods in Ecology and Evolution, 12(10), 1953–1968.

Loke, L. H., & Chisholm, R. A. (2023). Unveiling the transition from niche to dispersal assembly in ecology. Nature, 618(7965), 537–542.

Mahony, C. R., Cannon, A. J., Wang, T., & Aitken, S. N. (2017). A closer look at novel climates: New methods and insights at continental to landscape scales. Global Change Biology, 23(9), 3934–3955. 10.1111/gcb.13645

Mammola, S. (2019). Assessing similarity of nDdimensional hypervolumes: Which metric to use?. Journal of Biogeography, 46(9), 2012–2023.

Mammola, S. & Cardoso, P. (2020). Functional diversity metrics using kernel density n-dimensional hypervolumes. Methods in Ecology and Evolution, 11, 986–995.

McCann, K. S. & Gellner, G. (2020). Theoretical ecology: concepts and applications.

Menchetti, M., Guéguen, M. & Talavera, G. (2019). Spatio-temporal ecological niche modelling of multigenerational insect migrations. Proceedings of the Royal Society B, 286, 20191583.

Meireles, J. E., O’Meara, B., & Cavender-Bares, J. (2020). Linking leaf spectra to the plant tree of life. Remote sensing of plant biodiversity, 155–172.

Qiao, H., Escobar, L. E., Saupe, E. E., Ji, L., & Soberón, J. (2017). A cautionary note on the use of hypervolume kernel density estimators in ecological niche modelling. Global Ecology and Biogeography, 26(9), 1066–1070.

Pagel, M. (1999). Inferring the historical patterns of biological evolution. Nature, 401(6756), 877–884.

Peterson, A. T., Cobos, M. E., & JiménezDGarcía, D. (2018). Major challenges for correlational ecological niche model projections to future climate conditions. Annals of the New York Academy of Sciences, 1429(1), 66–77.

Pianka, E. R., Vitt, L. J., Pelegrin, N., Fitzgerald, D. B., & Winemiller, K. O. (2017). Toward a periodic table of niches, or exploring the lizard niche hypervolume. The American Naturalist, 190(5), 601–616.

Pili, A. N., Tingley, R., Sy, E. Y., Diesmos, M. L. L. & Diesmos, A. C. (2020). Niche shifts and environmental non-equilibrium undermine the usefulness of ecological niche models for invasion risk assessments. Scientific Reports, 10, 1–18.

R Core Team. (2024). R: A language and environment for statistical computing. Version 4.4.0. https://www.R-project.org/.

Revell, L. J. (2024). phytools 2.0: an updated R ecosystem for phylogenetic comparative methods (and other things). PeerJ, 12, e16505.

Schönland, S. (1918). A summary of the distribution of the genera of south african flowering plants: (With Special Reference to those found in the Divisions of Uitenhage And Port Elizabeth.). Transactions of the Royal Society of South Africa, 7(1), 19–58.

Singh, B. K., Delgado-Baquerizo, M., Egidi, E., Guirado, E., Leach, J. E., Liu, H., & Trivedi, P. (2023). Climate change impacts on plant pathogens, food security and paths forward. Nature Reviews Microbiology, 21(10), 640–656.

Soberón, J. (2007). Grinnellian and Eltonian niches and geographic distributions of species. Ecology letters 10, 1115–1123.

Soberón, J. M. (2010). Niche and area of distribution modeling: a population ecology perspective. Ecography 33, 159–167.

Soberón, J., & Arroyo-Peña, B. (2017). Are fundamental niches larger than the realized? Testing a 50-year-old prediction by Hutchinson. Plos one, 12(4), e0175138.

Soberón, J., & Peterson, A. T. (2020). What is the shape of the fundamental Grinnellian niche?. Theoretical Ecology, 13(1), 105–115.

Soberón, J. & Nakamura, M. (2009). Niches and distributional areas: concepts, methods, and assumptions. Proceedings of the National Academy of Sciences 106, 19644–19650.

Stull, G. W., Qu, X. J., Parins-Fukuchi, C., Yang, Y. Y., Yang, J. B., Yang, Z. Y., … & Yi, T. S. (2021). Gene duplications and phylogenomic conflict underlie major pulses of phenotypic evolution in gymnosperms. Nature Plants, 7(8), 1015–1025.

Tagliari, M. M., Bogoni, J. A., Blanco, G. D., Cruz, A. P., & Peroni, N. (2023). Disrupting a socio-ecological system: could traditional ecological knowledge be the key to preserving the Araucaria Forest in Brazil under climate change? Climatic Change, 176(2), 2.

Tingley, R., Vallinoto, M., Sequeira, F., & Kearney, M. R. (2014). Realized niche shift during a global biological invasion. Proceedings of the National Academy of Sciences, 111(28), 10233–10238.

Uyeda, J. C., & Harmon, L. J. (2014). A novel Bayesian method for inferring and interpreting the dynamics of adaptive landscapes from phylogenetic comparative data. Systematic biology, 63(6), 902–918.

Valladares, F. et al. (2014). The effects of phenotypic plasticity and local adaptation on forecasts of species range shifts under climate change. Ecology letters 17, 1351–1364.

Vilas, D., Fletcher Jr, R. J., Siders, Z. A., & Chagaris, D. (2022). Understanding the temporal dynamics of estimated environmental niche hypervolumes for marine fishes. Ecology and Evolution, 12(12), e9604.

Wadhwa, R.R., Williamson, D.F., Dhawan, A. and Scott, J.G. (2018). TDAstats: R pipeline for computing persistent homology in topological data analysis. Journal of open source software, 3(28), p.860.

Walter, H. (1979) Vegetation of the Earth and ecological Systems of the geo-biosphere. Second edition. Springer-Verlag New York Inc. 1979.

Wennekes, P. L., Rosindell, J. & Etienne, R. S. (2012). The neutral—niche debate: a philo-sophical perspective. Acta biotheoretica 60, 257–271.

Wiens, J. J. (2011). The niche, biogeography and species interactions. Philosophical Transactions of the Royal Society B: Biological Sciences, 366(1576), 2336–2350.

Winemiller, K. O., Fitzgerald, D. B., Bower, L. M. & Pianka, E. R. (2015). Functional traits, convergent evolution, and periodic tables of niches. Ecology letters, 18, 737–751.

Zhang, Z., Mammola, S., McLay, C. L., Capinha, C. & Yokota, M. (2020). To invade or not to invade? Exploring the niche-based processes underlying the failure of a biological invasion using the invasive Chinese mitten crab. Science of the Total Environment, 728, 138815.

