## Supplementary File for "The evolution of Hutchinsonian climatic niche hypervolumes in gymnosperms"

**TEXT S1**

**Adaptive shifts**

At the family level, the root value of the adaptive regime (θ) of the hypervolume volume was estimated as 22.22. At this level, 12 shifts were observed (Figure S3A), leading to (1) Taxaceae (θ = 21.470, PP [Posterior Probability] = 0.320), (2) Podocarpaceae (θ = 21.501, PP = 0.307), (3) Welwitchiaceae (θ = 17.977, PP = 0.577), (4) Gnetaceae (θ = 24.484, PP = 0.672), (5) Welwitchiaceae and Gnetaceae (θ = 18.550, PP = 0.636), (6) Gingkoaceae (θ = 16.252, PP = 0.818), (7) Calycanthaceae (θ = 20.241, PP = 0.544), (8) Amborellaceae (θ = 17.513, PP = 0.544), (9) Calycanthaceae and Amborellaceae (θ = 18.863, PP = 0.623), (10) Matoniaceae (θ = 19.225, PP = 0.605), (11) Psilotaceae (θ = 23.462, PP = 0.561), and (12) Matoniaceae and Psilotaceae (θ = 19.965, PP = 0.392). These results were almost identical across all three runs for each taxonomic level.

**Persistence diagrams**

We tested if the relative distinctiveness of the niche of each genus and family of gymnosperms evolved non-independently, by comparing the distance between the centre of mass of the niche hypervolume of each genus or family to the centre of mass of the joint hypervolume of all gymnosperms (i.e., the "centroids") (Figures S2E, S3F). Greater distance would indicate that a given genus or family have evolved to exploit unique niches relative to the average niches of all gymnosperms. If the ability to exploit such niche boundary conditions are shared among closely related species, then our centroid analysis should reveal signs of non-independent evolution. However, our results showed no evidence of phylogenetic signal for the centroid distances at the genus or family levels (Table S3).

**SUPPLEMENTARY TABLES AND FIGURES**

**Table S1.** Families and genera used in the study with the respective DOI numbers from GBIF.

| **Downloaded taxon** | **GBIF DOI** | **Families included** | **Genera included** |
| --- | --- | --- | --- |
| Cycadopsida | 10.15468/dl.qjgjhg | Cycadaceae | *Cycas* |
|  |  | Zamiaceae | *Bowenia* |
|  |  |  | *Ceratozamia* |
|  |  |  | *Dioon* |
|  |  |  | *Encephalartos* |
|  |  |  | *Lepidozamia* |
|  |  |  | *Macrozamia* |
|  |  |  | *Microcycas* |
|  |  |  | *Stangeria* |
|  |  |  | *Zamia* |
| Ginkgoopsida | 10.15468/dl.zxemsp | Ginkgoaceae | *Ginkgo* |
| Gnetopsida | 10.15468/dl.5hykeh | Ephedraceae | *Ephedra* |
|  |  | Gnetaceae | *Gnetum* |
|  |  | Welwitschiaceae | *Welwitschia* |
| Pinopsida | 10.15468/dl.r34zkh | Araucariaceae | *Agathis* |
|  |  |  | *Araucaria* |
|  |  | Cupressaceae | *Athrotaxis* |
|  |  |  | *Austrocedrus* |
|  |  |  | *Callitris* |
|  |  |  | *Calocedrus* |
|  |  |  | *Cryptomeria* |
|  |  |  | *Cunninghamia* |
|  |  |  | *Cupressus* |
|  |  |  | *Diselma* |
|  |  |  | *Fitzroya* |
|  |  |  | *Juniperus* |
|  |  |  | *Libocedrus* |
|  |  |  | *Metasequoia* |
|  |  |  | *Microbiota* |
|  |  |  | *Pilgerodendron* |
|  |  |  | *Platycladus* |
|  |  |  | *Sequoia* |
|  |  |  | *Sequoiadendron* |
|  |  |  | *Taiwania* |
|  |  |  | *Taxodium* |
|  |  |  | *Tetraclinis* |
|  |  |  | *Thuja* |
|  |  |  | *Thujopsis* |
|  |  |  | *Widdringtonia* |
|  |  | Pinaceae | *Abies* |
|  |  |  | *Cedrus* |
|  |  |  | *Keteleeria* |
|  |  |  | *Larix* |
|  |  |  | *Picea* |
|  |  |  | *Pinus* |
|  |  |  | *Pseudotsuga* |
|  |  |  | *Tsuga* |
|  |  | Podocarpaceae | *Afrocarpus* |
|  |  |  | *Dacrycarpus* |
|  |  |  | *Dacrydium* |
|  |  |  | *Halocarpus* |
|  |  |  | *Lepidothamnus* |
|  |  |  | *Manoao* |
|  |  |  | *Microcachrys* |
|  |  |  | *Nageia* |
|  |  |  | *Pherosphaera* |
|  |  |  | *Phyllocladus* |
|  |  |  | *Podocarpus* |
|  |  |  | *Prumnopitys* |
|  |  |  | *Retrophyllum* |
|  |  |  | *Saxegothaea* |
|  |  | Sciadopityaceae | *Sciadopitys* |
|  |  | Taxaceae | *Cephalotaxus* |
|  |  |  | *Taxus* |
|  |  |  | *Torreya* |
| Amborellaceae (outgroup) | 10.15468/dl.88h5pn | Amborellaceae | *Amborella* |
| Calycanthaceae (outgroup) | 10.15468/dl.k732vq | Calycanthaceae | *Calycanthus* |
| Matoniaceae (outgroup) | 10.15468/dl.htvb4x | Matoniaceae | *Matonia* |
| Psilotaceae (outgroup) | 10.15468/dl.pyugx9 | Psilotaceae | *Psilotum* |
|  |  |  | *Tmesipteris* |

**Table S2.** Phylogenetic signal using Blomberg's *K* (Blomberg et al., 2003) and Pagel's λ (Pagel, 1999) of hypervolume volume and holes at the family level.

| **Trait** | 𝝺 | **logL** | **logL0** | **p** | **K** | **p** |
| --- | --- | --- | --- | --- | --- | --- |
| Hypervolume | 0.000 | -35.979 | -35.979 | 1.000 | 0.776 | 0.626 |
| Mean distance | 0.000 | 4.933 | 4.933 | 1.000 | 0.843 | 0.440 |
| Maximum distance | 0.000 | -7.764 | -7.764 | 1.000 | 0.737 | 0.773 |
| SD distance | 0.000 | -4.220 | -4.220 | 1.000 | 0.702 | 0.863 |

**Table S3.** Phylogenetic signal using Blomberg's K (Blomberg et al., 2003) and Pagel's 𝝺 (Pagel, 1999) of PCA centroids at the genus and family level.

| **Level** | 𝝺 | **logL** | **logL0** | **p** | **K** | **p** |
| --- | --- | --- | --- | --- | --- | --- |
| Genus | 0 | -66.141 | -66.14 | 1 | 0.493 | 0.169 |
| Family | 2.108 | -11.457 | -12.968 | 0.082 | 1.003 | 0.2 |

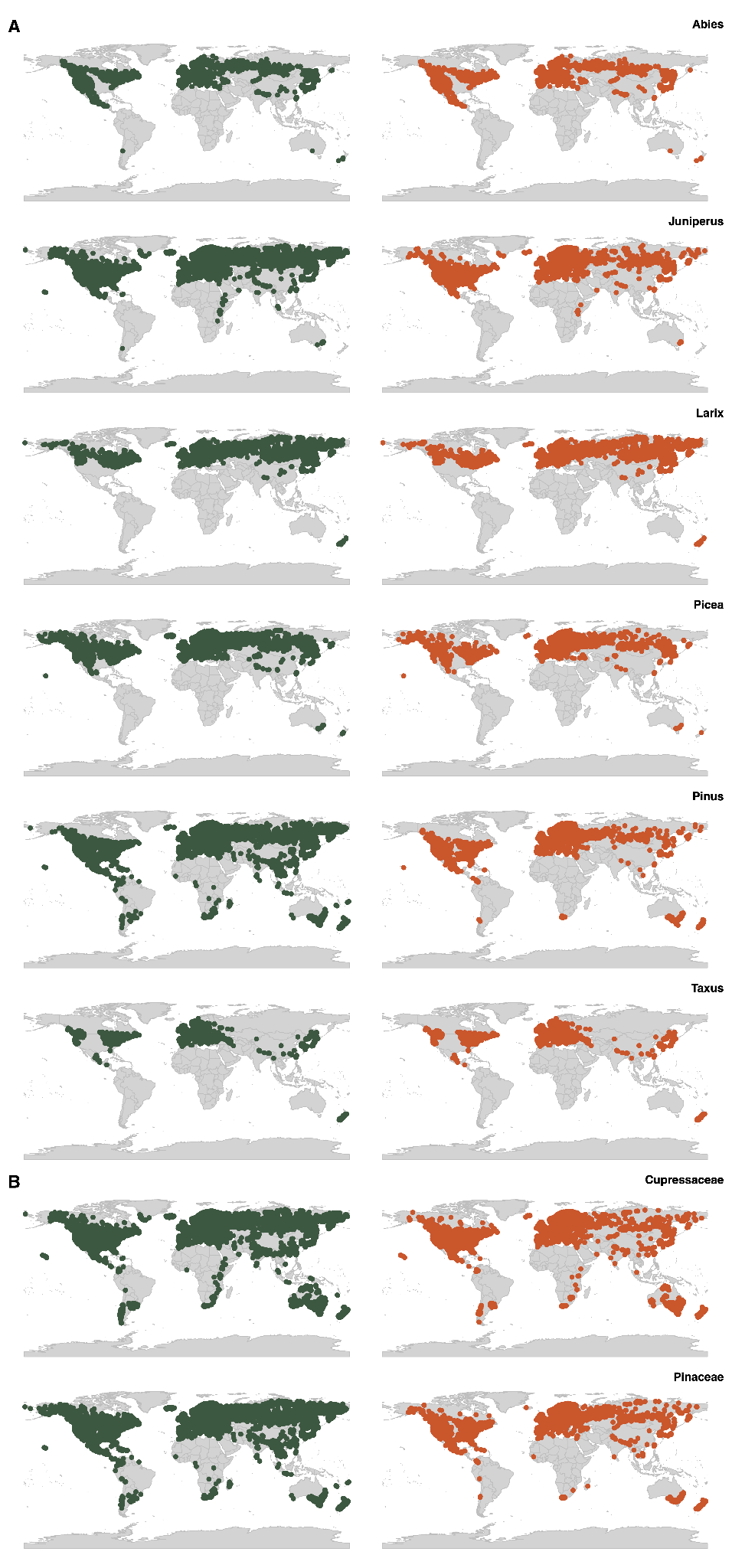

**Figure S1.** Occurrence points for the genera and families with more than 50,000 data points. At the left (green), are all available occurrence points in GBIF. At the right (orange), the subset of 50,000 data points for this study is shown. **A.** Occurrence points at the genus level. **B.** Occurrence points at the family level.

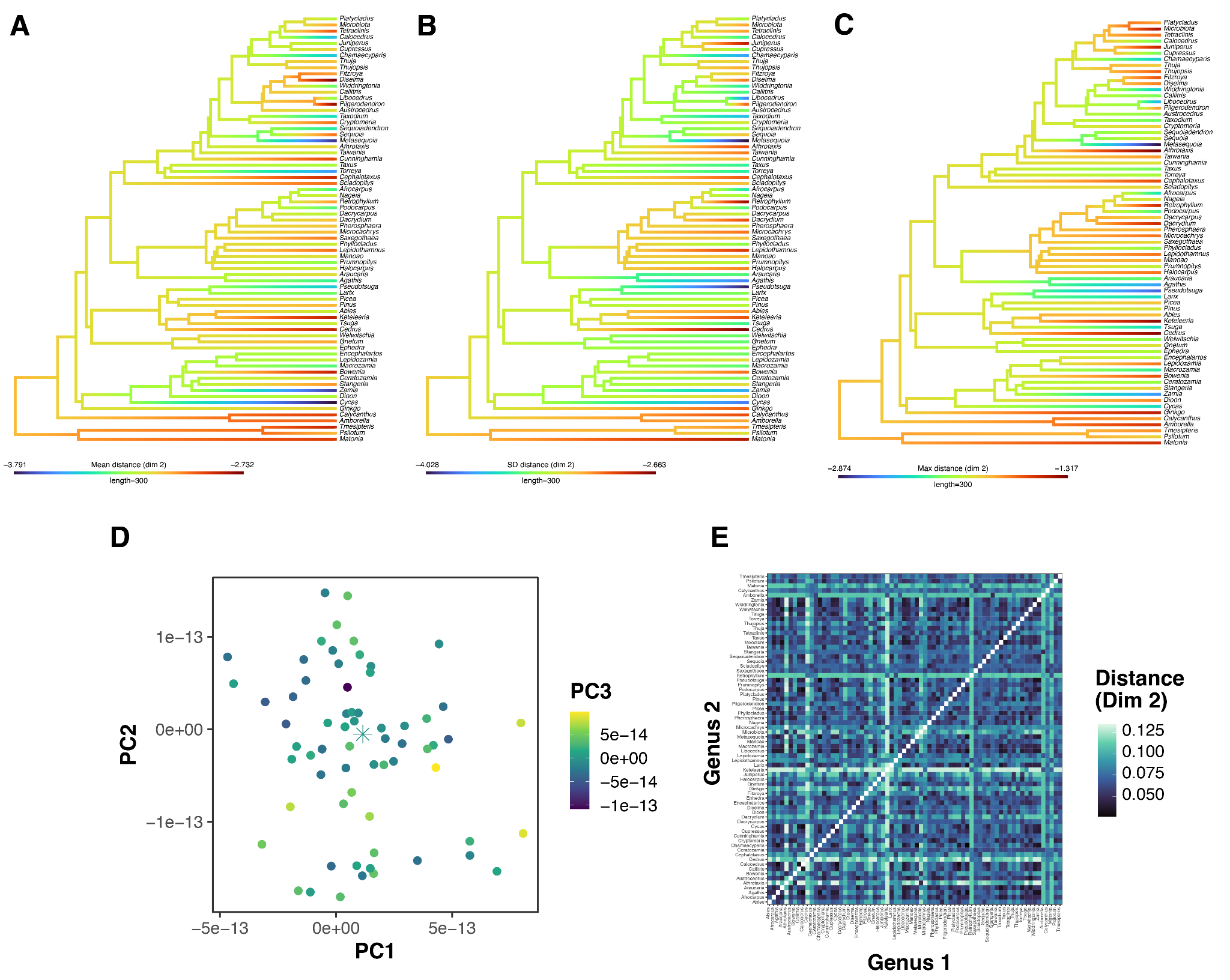

**Figure S2.** Phylogenetic signal and centroid analysis at the genus level. **A.** Character mapping of mean distance of points at dimension 2. **B.** Character mapping of the standard deviation of distance of points at dimension 2. **C.** Character mapping of the maximum distance of points at dimension 2. **D.** Centroids of the PCA analysis for each genus. Circles represent each genus centroid, while the star represents the centroid for all genera. **E.** Bottleneck pairwise distance of persistence diagrams. Darker values represent closer bottleneck distances.

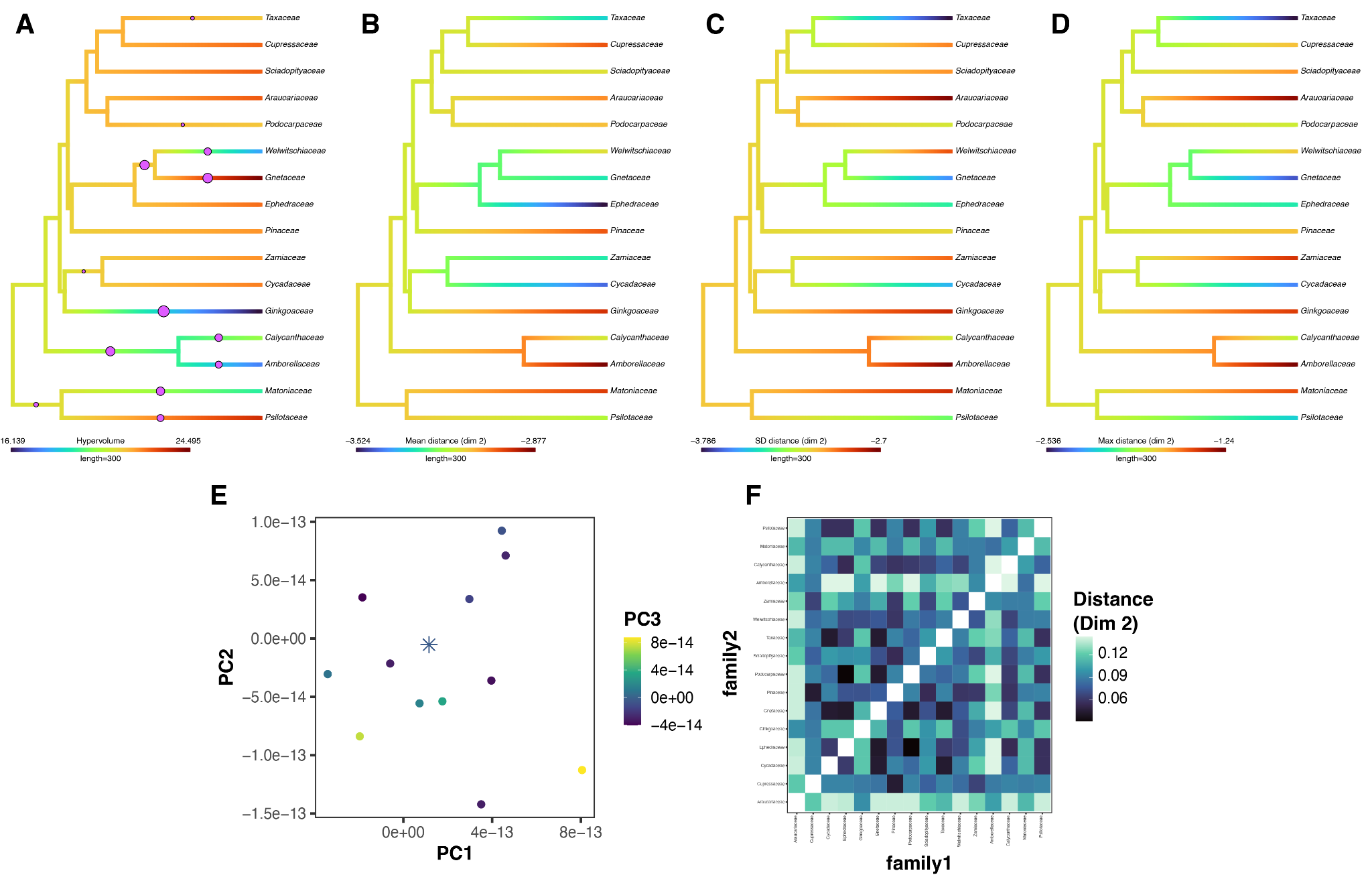

**Figure S3**. Phylogenetic signal, adaptive shifts, and centroid analysis at the genus level. **A.** Character mapping of the hypervolume volume. Circles represent the adaptive shifts with a posterior probability higher than 0.3. **B.** Character mapping of mean distance of points at dimension 2. **C.** Character mapping of the standard deviation of distance of points at dimension 2. **D.** Character mapping of the maximum distance of points at dimension 2. **E.** Centroids of the PCA analysis for each family. Circles represent each family centroid, while the star represents the centroid for all families. **F.** Bottleneck pairwise distance of persistence diagrams. Darker values represent closer bottleneck distances.

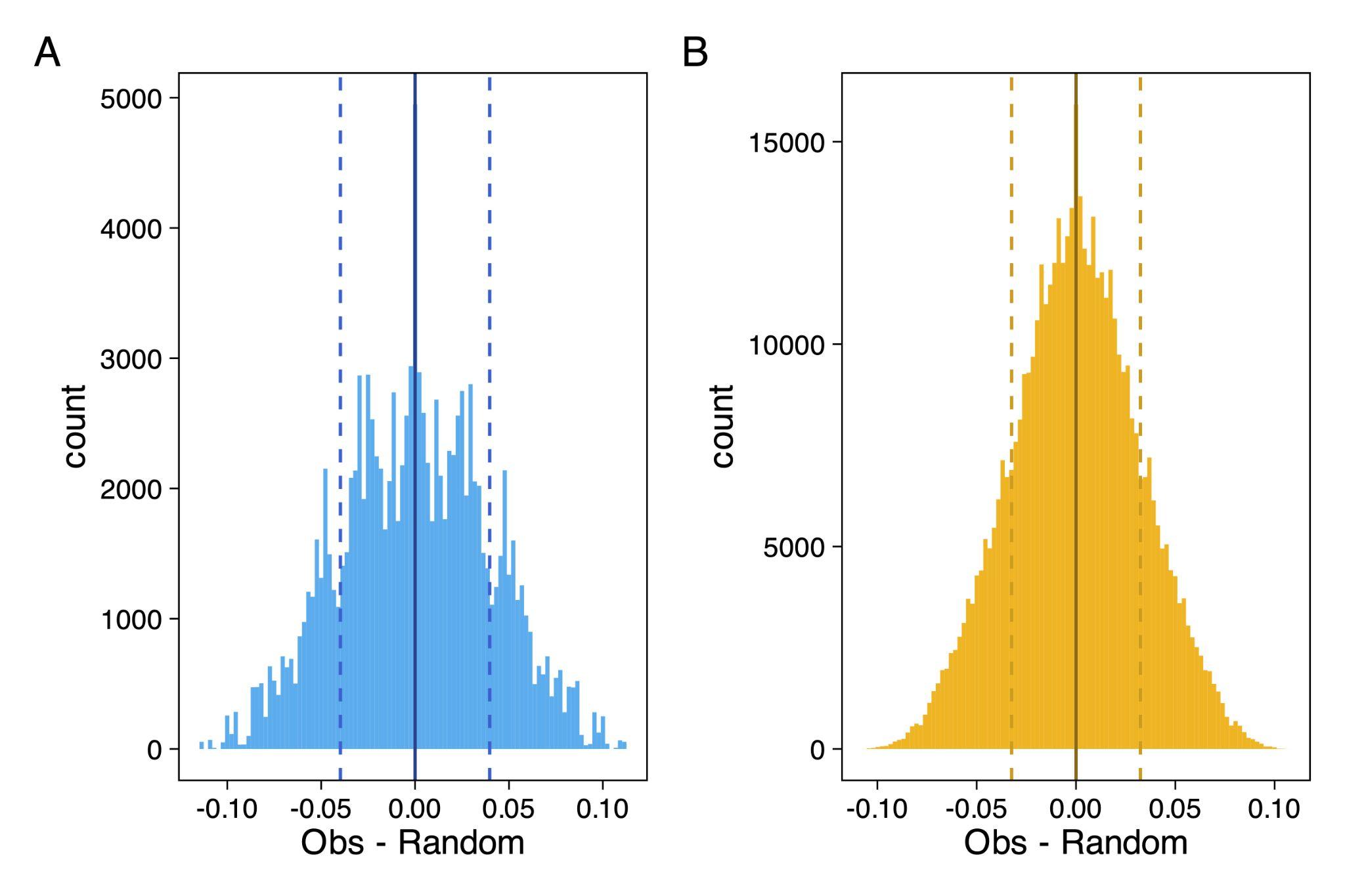

**Figure S4.** Differences between the pairwise distance matrix calculated with the empirical data and a random pairwise matrix. **A.** Family level analysis**. B.** Genus level analysis. Solid line = mean, dashed = mean + or – 1 standard deviation
